# Forecasting Carbon Dioxide Emissions Of Bahrain Using Singular Spectrum Analysis And ARIMA Hybrid Model

**DOI:** 10.1101/2025.03.26.645400

**Authors:** Zahrah Fayez Althobaiti

## Abstract

Predicting the economic implications of gas emissions and their repercussions is critical to policymakers, especially given the current increasing trend in volume. Therefore, study on gas emission prediction is required. A hybrid model is proposed for forecasting CO_2_ emissions of Bahrain (BH) in this study. Singular Spectrum Analysis (SSA) and Auto Regressive Integrated Moving Average are the two techniques that make up the hybrid model (ARIMA). In this model, the time series of CO_2_ emissions are first divided into a number of sub-series corresponding to some tendentious and oscillation (periodic or quasi-periodic) components and noise by using SSA. Each sub-series is then predicted individually through an appropriate ARIMA model. Finally, a correction procedure is carried out for the sum of the prediction results to ensure that the superposed residual is a pure random series. The result of forecasting obtained by using hybrid SSA-ARIMA was compared with ARIMA to assess its superiority. While, the application of SSA was to improve the accuracy of forecasting of CO_2_ emissions in BH. The data used in this study are the annual time series data in three different periods from 1990-2018, 2000-2018 and 2003-2018 for CO_2_ emissions in BH. The Mean Absolute Percentage Error (MAPE) and Root Mean Square Error (RMSE) criteria are used to assess each forecasting method’s level of forecasting accuracy. The results indicated that the SSA-ARIMA method is superior to the ARIMA model as the best forecasting method for CO_2_ emis-sions.

## 1 Introduction

One of the most embarrassing and explosive issues today, particularly in developing nations, is global warming and climate change((Destek and Sarkodie, 2019); (Khalid *et al*., 2021)).The industrialization and urbanization processes that have accelerated recently are to blame for the rise in greenhouse gas (GHG) emissions. One of the main causes of GHGs and a primary driver of global warming is carbon dioxide (CO_2_) (Yang and Usman, 2021). Accordingly, ((Hossain *et al*., 2017); (Bonga and Chirowa, 2014)) accentuated that human activities are one of the main causes of carbon dioxide emissions, with the production of energy from coal, oil, and natural gas as well as the use of petroleum goods for travel, aviation, and automobile trips being the most significant.

Bahrain is one of the Arab Gulf countries (AGC), whose oil and gas constitute the main component of its economy. Bahrain is the first country to discover oil in the Arabian Gulf, which had a great impact on changing all aspects of life for the Gulf countries, as well as the country’s ability to pump large investments ((Qader, 2009); (Yang and Usman, 2021); (Yang *et al*., 2021)). In this regard, the emission of CO_2_ gas resulting from the use of energy is one of the negative effects on the environment, thus the most important reasons for forecasting CO_2_ emission in Bahrain. However, in order to predict CO_2_ emissions, one must extrapolate time series values from observed time series, since it has been demonstrated to help increase public awareness of how to prevent environmental diffi-culties The prediction of CO_2_ emissions has grown to be a concern on a global scale (Abdullah and Pauzi, 2015). Therefore, a better comprehension of Bahrain’s historical CO_2_ emission is crucial to developing a reasonable forecast of the country’s future CO_2_ emissions.

Forecasting is significant in both academics and industry, as it is a key component of strategic, tactical, and operational activities. Combining predictions is a well-established approach for improving forecasting accuracy that takes advantage of different information and computational resources (Bunn, 1996), which dates back to Bates and Granger’s foundational work 37 years ago (Bates and Granger, 1969). They demonstrated that an appropriate linear combination of two distinct time series forecasts can result in a superior forecast than the two original forecasts. Many additional experts, Clemen, (1989) and Chan *et al*. (1999) have published empirical results that suggest that combining forecasts outperforms individual forecasting methods.

Recently, multiple sectors have employed the Singular Spectrum Analysis (SSA) approach for forecasting. SSA is a recently developed non-parametric data-driven method for modelling noisy, non-stationary, and/or non-linear time series (Hassani, 2007). The SSA decomposition of the original time series into the sum of independent components can be used to illustrate the trend, oscillatory activity (periodic or quasiperiodic components), and noise (Golyandina *et al*., 2001). The fact that SSA only needs two parameters to model the time series under consideration, set it apart from other non-parametric methods in a big way (Hassani and Zhigljavsky, 2009). In recent years, SSA has proven to be a valuable tool for assessing and forecasting time series variables such as variations, periodicities, and trends ((Ghil *et al*., 2002); (Hassani *et al*., 2009); (Kumar, 2015)). In the recent studies of future prediction of time series, SSA has shown to be more accurate than numerous well-known approaches (Zhang *et al*., 2011). The Death series is an example of a seemingly complex series with potential structure that can be simply analysed using SSA and could serve as a good illustration of how SSA can be used successfully (Hassani, 2007; Hassani *et al*., 2010). Also, when employed on data with a complicated structure, SSA provides superior forecasting results than other methods (Z *et al*., 2011).

Consequently, SSA hybrid modelling, as well as other approaches, is widely used (Li *et al*., 2014). The hybrid technique works by combining two or more models to provide more accurate forecasts or prediction outcomes (Vahabie *et al*., 2007). SSA is a pre-processing approach that can be used in conjunction with other methods (Li *et al*., 2014). As superiority to conventional statistical models, the grey model based on the fractional calculus becomes a hot topic and show great potentials with excellent performance. In this study hybrid SSA-ARIMA method was employed to predict CO_2_ emissions in BH, and compared with ARIMA method.

The prediction using ARIMA models statistical method is usually viewed as providing more accurate predictions than econometric methodologies (Song *et al*., 2003). Also, in terms of forecasting performance, ARIMA models outperformed the multivariate models (Du Preez and Witt, 2003). Moreover, ARIMA models outperform naive models and smoothing approaches in terms of overall performance (Goh and Law, 2002). ARIMA models were created in the 1970s by Box and Jenkins, and its identification, estimation, and diagnostics method is based on the notion of parsimony (Asteriou and Hall, 2015). That is; when the original time series is not stationary, the first order difference process Δ*Y* or second order differences Δ^2^*Y*, and so on, can be investigated. While, If the differenced process is a stationary process, ARIMA model of that differenced process can be found in practice if differencing is applied, usually d=1, or maybe d=2, is enough. When fitting time series, the ARIMA model has the benefit of being able to employ a combination of auto regression, difference, and moving average of different orders to generate the ARIMA (p, d, q) model, which can convey multiple types of information of time series. By choosing the right parameters, accurate forecasts can be made. The ARIMA model, on the other hand, demands that the sequence be steady after differentiation. At the moment, ARIMA is still mostly utilised for comparisons with other sophisticated models.

Accordingly, the combination of several models will greatly improve the reliability of prediction. The advantage-seeking combination model had the characteristic of avoiding drawbacks while pursuing advantages. The benefits of a single model might be obtained while the problems could be avoided by combining both models. This is also why we must develop combinatorial models in order to get smaller relative errors. The combined forecasting modelling methodology combines two or more predictive methodologies to predict the same prediction problem. The combination forecast, which is based on theoretical research and real application, maximizes the benefits of each forecasting model while also allowing for future forecast conditions to evolve. It has the potential to improve forecast accuracy while also increasing prediction stability (Tian et al., 2012).

This study thus aimed to combine prediction data from a number of mixed time series forecasting models to predict CO_2_ emissions in BH using the hybrid SSA-ARIMA method, and compared with ARIMA model. The findings will serve as a guide for future energy planning as well as the overall economic improvement of Bahrain (BH).

## 2 Literature Review

Nyoni & Bonga, (2019) used the Box-Jenkins ARIMA technique to model and predict CO_2_ using annual time series data on CO_2_ emissions in India from 1960 to 2017. Test results show that India’s CO_2_ emission statistics is incorrect (2). According to the study, the ARIMA (2, 2, 0) model, with an estimated 3.89 million kt of CO_2_ by 2025, has the best match for predicting India’s annual CO_2_ emissions for the following 13 years.

In Bangladesh, Hossain et al. (2017) created acceptable statistical models for projecting CO_2_ emissions from 1972 to 2013. Annual CO_2_ emissions statistics in metric tons per capita (kt.), Gaseous Fuel Consumption (GFC), Liquid Fuel Consumption (LFC), and Solid Fuel Consumption (SFC) data were obtained from the World Bank data base (SFC). To develop Autoregressive Integrated Moving Average (ARIMA) models, the Kwiatkowski Phillips Schmidt Shin (KPSS) test was utilized to determine the unit root property of the research variables. The models employed the Akaike Information Criterion (AIC), Schwarz Information Criterion (SIC), Coefficient of Determination (R2), Root Mean Square Error (RMSE), Mean Absolute Error (MAE), and Bias Proportion (BP) measures. The ARIMA models outperformed the Holt-Winters Non Seasonal (HWNS), Smoothing, and Artificial Neural Networks (ANN) models in static forecasting from 1972 to 2013. In addition, ARIMA models were employed to forecast data from 2011 to 2013, and the Mean Absolute Percentage of Error (MAPE) for GFC, LFC, and SFC were 2.8, 8.4, and 12.9 kt, respectively. In the Chow forecast test, the stability of the forecasting process is further examined using real-world data. ARIMA also did out-of-sample static forecasting from 2014 to 2025, and the expected values for GFC, LFC, and SFC in 2025 are 53034.79, 15926.17, and 9579.49 kt, respectively, which is alarming. Thus, Bangladesh may be able to make better appropriate climate policies because of its CO_2_ emissions from gasoline used.

Also, studies by Mahmoudvand et al., (2015) shows that in most circumstances the forecasting accuracy of the SSA approach is higher than the forecasting accuracy of the Hyndman and Ullah method. The RSSA performs marginally better than the VSSA when comparing two SSA techniques of RSSA and VSSA. Aside from that, the basic methods for selecting parameters in the forecasting are similar to those used in the reconstruction stage. The reconstruction stability can be even more important than the reconstruction accuracy for forecasting, which is a significant difference. Furthermore, past simulation and theory studies demonstrated that selecting a window length L less than half the length of the time series N is preferable. N/3 is one of the suggested values (Golyandina, 2010). Aside from that, theoretical breakthroughs and research findings reinforced the SSA forecasting potential in economic and financial time series. The SSA can really produce similar and better results than other forecasting methods (Hassani and Thomakos, 2010). As a result, it is evident that SSA forecasting offers a lot of potential in terms of obtaining more accurate forecasting data.

Moreover, Hassani, (2007) used Singular Spectrum Analysis (SSA) methodologies to evaluate monthly accidental death time series in the United States and compared them to existing predicting procedures. The results demonstrated that SSA was more accurate. The author used SSA to decompose a time series into three separate and interpretable components: slowly shifting trend, oscillatory components, and structureless noise. Between 1973 and 1978, SSA was employed to a time series of monthly accidental deaths in the United States. In terms of mean absolute error (MAE) and mean relative absolute error (MRAE), experimental results indicated a superior forecasting method, demonstrating a typical example of SSA successful use. The author recommended low-frequency boundary selection and threshold determination before finishing trend extraction using time series spectral analysis. The suggested methodology’s ease of use was demonstrated through simulations utilizing unemployment data time series from Alaska.

Hassani and Thomakos, (2010) examined SSA’s theoretical and analytical method for economic and financial time series forecasting in the same direction. The study provided two novel SSA versions, as well as a range of simulation findings and applications, in addition to existing forecasting methodologies. The authors investigated SSA in the context of economic and financial time series, giving a broad overview of the method as well as forecasting skills for individual time series components or rebuilt time series. The calculation of window length L and eigentriple number r for time series reconstruction, as well as SSA forecasting, has received a lot of attention. The outcomes backed up SSA’s promise in predicting economic and financial time series, as did the theoretical advances.

Another SSA application investigated into the feasibility of using SSA to predict mortality in nine European countries -Belgium, Denmark, Finland, France, Italy, Netherlands, Norway, Sweden, and Switzerland (Mahmoudvand *et al*., 2015). The Hyndman-Ullah predicting model was used to compare and assess the SSA’s forecasting skills. The reason the authors picked the suggested methodology was that they believed SSA recog-nised important structural characteristics including time series trend, oscillatory components, and noise and dissected time series into a subset of distinct time series components. Based on empirical considerations, the length of the window L and the quantity of eigen-triples utilised for time series reconstruction were selected, and the results were verified using actual data.

The research by Liu et al. (2015) proposed a hybrid approach for estimating medium and long-term software failure times. The hybrid model is composed of Singular Spectrum Analysis (SSA) and Auto Regressive Integrated Moving Average (ARIMA). In this model, the time series of software failure time are first divided into several subseries using SSA, each of which corresponds to some tendentious and oscillation (periodic or quasi-periodic) components and noise. Each subseries is then predicted using an appropriate ARIMA model, and finally the sum of the prediction results is corrected to ensure that the superposed residual is a pure random series. The software failure data from two actual projects is reviewed as case studies. The outcomes were contrasted with forecasts made using ARIMA and Singular Spectrum Analysis-Linear Recurrent Formulae (SSA-LRF). The hybrid model operates most effectively.

Moreover, Hassani et al. (2015) investigated into the possible benefits of utilizing SSA to forecast tourism demand. The authors used monthly data on visitor arrivals in the United States from 1996 to 2012 to analyse the SSA predicting performance. SSA predictions were compared to those from ARIMA, exponential smoothing, and neural networks, among other forecasting methods. The findings suggested that the SSA method produced projections that performed much better (statistically) than other approaches. The main finding was that the SSA has significant benefits in projecting visitor arrivals in the United States and should be considered for additional tourism demand forecasting research.

## 3 Research Methodology

In this study, the hybrid SSA-ARIMA and ARIMA were utilized to predict CO_2_ emission in BH from 1990 – 2018, 2000-2018, and 2003-2018 as presented below.

### 3.1 Singular Spectrum Analysis (SSA)

SSA decomposes a time series into independent elements that describe the trend, oscillatory conduct (cyclic or quasi-cyclic elements), and noise. Several of these elements are picked and preserved for the predicting purpose. SSA has been effectively applied in a variety of fields, including hydrology (Vautard *et al*., 1992), geophysics, and others (Vautard and Ghil, 1989).

Let *y* = [*y*_1_, *y*_2_,…, *y_N_*] be a time series of length N. Decomposition and reconstruction are the two stages of the SSA approach (Golyandina *et al*., 2001).

#### Decomposition

Embedding and Singular Value Decomposition are the two steps in this phase.

#### Embedding

The definition of a trajectory matrix for the initial time series y is the main outcome of this phase. A windows length L (L=N/2, to be defined by the user) is associated with the matrix. Let K=N-L+1 be the trajectory matrix, and the trajectory matrix is defined as:

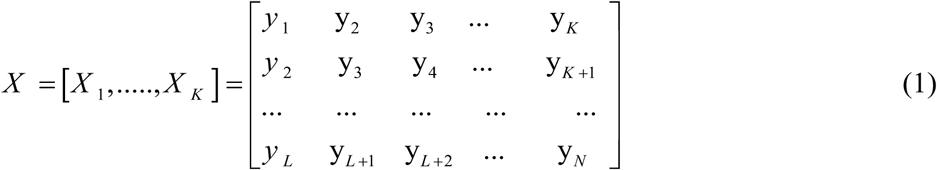

All of the elements along the diagonal i+j=const are equal in the trajectory matrix, which is a Hankel matrix. And let *S* = *XX^T^*, this call S the covariance matrix.

#### Singular Value Decomposition (SVD)

From X, define the covariance matrix *S* = *XX^T^*. The SVD of *XX^T^* creates a set of L. *λ*_1_≥*λ*_2_≥…≥*λ_L_* eigenvalues and *U*_1_,*U*_2_,..,*U_L_* eigenvectors (commonly referred to as Empirical Orthogonal Functions (EOF)). The trajectory matrix’s SVD can then be rep-resented as:

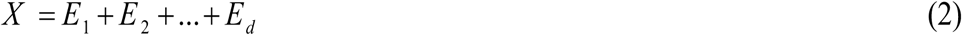

Where; 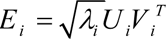, d is the rank of *XX^T^* (that is the number of non-zero eigenvalues) and *V*_1_,*V*_2_,..,*V_d_* are the Principal Components (PC), defined as 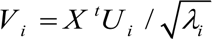 The i-th eigentriple of the matrix X is represented by the collection (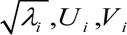)

#### Reconstruction

There are two processes in the reconstruction stage: grouping and averaging.

#### Grouping

In the grouping phase, the SVD terms in the trajectory matrix’s SVD are grouped. The purpose of grouping SVD components is to express the trajectory matrix of the original time series as a sum of trajectory matrices that uniquely specify the time series’ expansion into a total of additive components. The user chooses r out of d eigentriples in this stage. Let *I*={i1.i2,…,ir} be a group of r with (1≤*r*≤*d*) chosen eigentriples and *X_I_* = *X*_*i*1_ + *X*_*i*2_ + … + *X*_*ir*_. *X_I_* is connected to the “signal” of y while the rest of the (d-r) eigentriples illustrate the error term ε.

#### Averaging

The deterministic components of the time series are then reconstructed using the group of r elements chosen in the former stage. The main idea is to use the Hankelization technique to transform each terms *X_I_* = *X*_*i*1_ + *X*_*i*2_ + … + *X*_*ir*_ into a rebuilt time series *y*_*i*1_, *y*_*i*2_,…, *y_ir_*. H(), also known as diagonal averaging, is a function that calculates the average of if *i* + *j* = *k* + 1 and *z_ij_* is an element of a generic matrix Z, then averaging *z_ij_* yields the k-th term of the reconstructed time series. That is, H(Z) is an N-length time series rebuilt from matrix Z. The rebuilt time series approximates y at the end of the av-eraging phase.

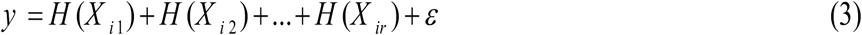

#### Parameter selection

SSA just requires two parameters, as previously stated: the window length L and the number of components r. Data from the time series under inquiry or other indices could be used to calculate L and r values.

- **The window length L**: The only parameter in the decomposition stage is the window length L. The challenge at hand and preliminary data regarding the time series must be taken into consideration while choosing the appropriate window length (Hassani and Zhigljavsky, 2009). Knowing that the time series may have a periodic component with an integer period, it is advised to select the window length proportional to that period to improve the separability of these components. Theoretical findings indicate that L should be sufficiently large but not greater than N /2 (Hassani and Thomakos, 2010).
- **The number of components r**: The definition of r is based on the principle of separability, or how well the elements can be separated. The following quantity (called the weighted correlation or w-correlation) is a natural measure of the dependence between two time series. Let *Y_T_*^(1)^ and *Y_T_*^(2)^ be two reconstructed time series;

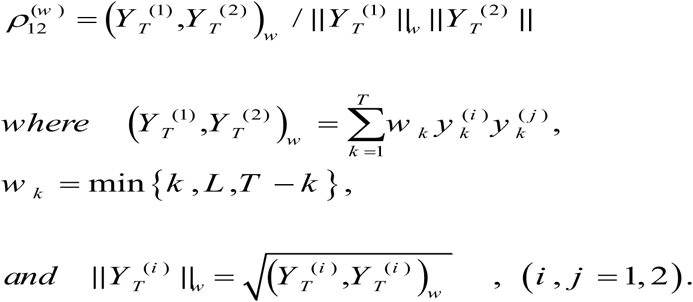

Two reconstructed components are said to be well separated if they have zero w-correlation. The components should be grouped together and may even belong to the same component in the SSA decomposition, according to large values of w-correlations between the reconstructed components (Alexandrov and Golyandina, 2005).

In equation (2), the norm of X is most positively affected by the first elementary matrix E1 with the norm 1, and it is least positively affected by the last elementary matrix Ed with the norm d. The decision of where to truncate the summation of Equ (2) in order to construct a decent approximation of the original matrix depends on the location of the truncation. The plot of the eigenvalues 1,…,, d provides an overall picture of the values of the eigenvalues. The noise component of the series is often represented by a progressively diminishing succession of eigenvalues. The identification of the eigentriples that belong to the same harmonic component of the series is made possible by similar eigen-value values. Selecting the groups can also benefit from the periodogram analysis of the original time series. The harmonic components of the series are linked to sharp sparks in the graph.

#### SSA forecasting

There are various researches describe about this forecasting algorithm. From this study, the forecasting algorithms are defined in ((Golyandina *et al*., 2001), (Hassani and Thomakos, 2010),(Hassani *et al*., 2009)), is as follows:

a. Let consider the time series, *Y_N_* = (*y*_1_,…*y_N_*) with length N
b. Assume window length L, 1 < L < N.
c. Create the time series’ trajectory matrix *Y_N_* in equ (1)
d. Construct eigenvectors *U*1, …,*UL* from the SVD of *X*
e. Estimate matrix

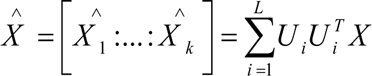
f. Hankellization step: construct matrix in equ (3)
g. Set 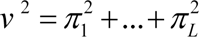 where *π_i_* is the last component of the vector *Ui* (*i* = 1, …,*L*) where *v*^2^ < 1
h. Determine vector *A* = (*α*_1_,…*α*_*L*−1_).

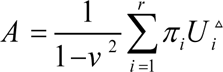

Assume that *π_i_* is the last component of the vector *Ui* (= 1, …, *L*) and 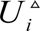*^]^* is the vectors of the first L – 1 component of the eigenvectors *Ui* where:

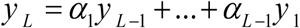

(g) For predicting procedure, define the time series by the formulas:

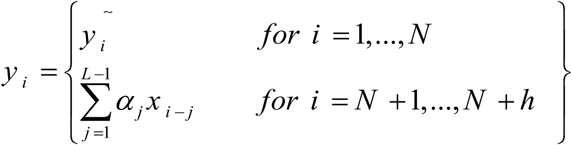

The numbers of *^y N^* ^+1,…, *y*^ *^N^* ^+^*^h^* are from the h terms of the SSA recurrent forecast.

### 3.2 ARIMA model

The ARIMA model is one of the most widely used statistical models for time series forecasts (Box and Jenkins, 1976). Its forecast principle is to transfer a non-stationary time series into a stationary time series first. As a result, the dependent variable will be described as a model that only yields its lag value, as well as the actual and lag values of the random error term. The following are the steps in the prediction phase: (Wang *et al*., 2018).

**Phase 1:** Smooth the time data with a differential tool. In theory, stationarity ensures that the fitted curve formed by sampling time series can continue inertially along the present form in the future, i.e., the data’s mean and variance should not be significantly changed.

**Phase 2:** Create a model that is autoregressive (AR). The autoregressive model is a way of forecasting itself using the variable’s historical result data, and it illustrates the link between current value and previous value. It has the following formula:

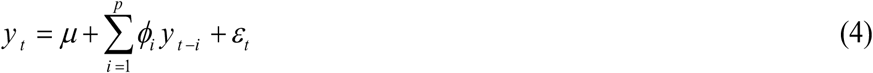

Where *^yt^* represents the current value, *µ* indicates the constant term, *p* denotes the order, *ϕ_i_* is the autocorrelation coefficient, and *ε_t_* represents the error.

**Phase 3:** Create a model based on moving averages (MA). In the autoregressive model, the moving average model concentrates on the accumulation of error components. Random fluctuations in forecasts can be successfully eliminated. It has the following for-mula:

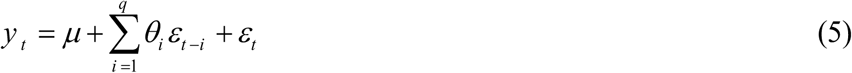

Where *θ_i_* is the MA formula’s correlation coefficient.

**Phase 4:** Create an autoregressive moving average model by combining AR and MA (ARMA). The following is the exact formula. The orders of the autoregressive and moving average models, respectively, are p and q in this formula. The correlation coefficients of the two models, *ϕ_i_* and *θ_i_*, respectively, must be solved.

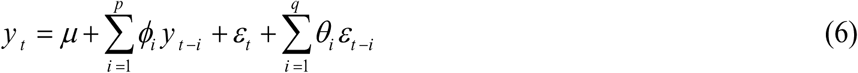

### 3.3 Analysis of hybrid SSA-ARIMA model

The SSA-ARIMA hybrid model uses sub-series components (RCs) produced by SSA on original data to forecast the annual time series of CO2 emissions in SA. The following steps substantially summarize the methodological steps for the SSA-ARIMA model.

1. SSA first divides the original time series into principal components (PCs).
2. A new series for each variable is created by summing the appropriate RCs, which are chosen based on the trend or period of each series.
3. The ARIMA models are used to forecast the new series of RCs, respectively.
4. Sum all results of the forecast to get predicted of a new time series.

This procedure is described in Figure 2.

**Figure 1.**
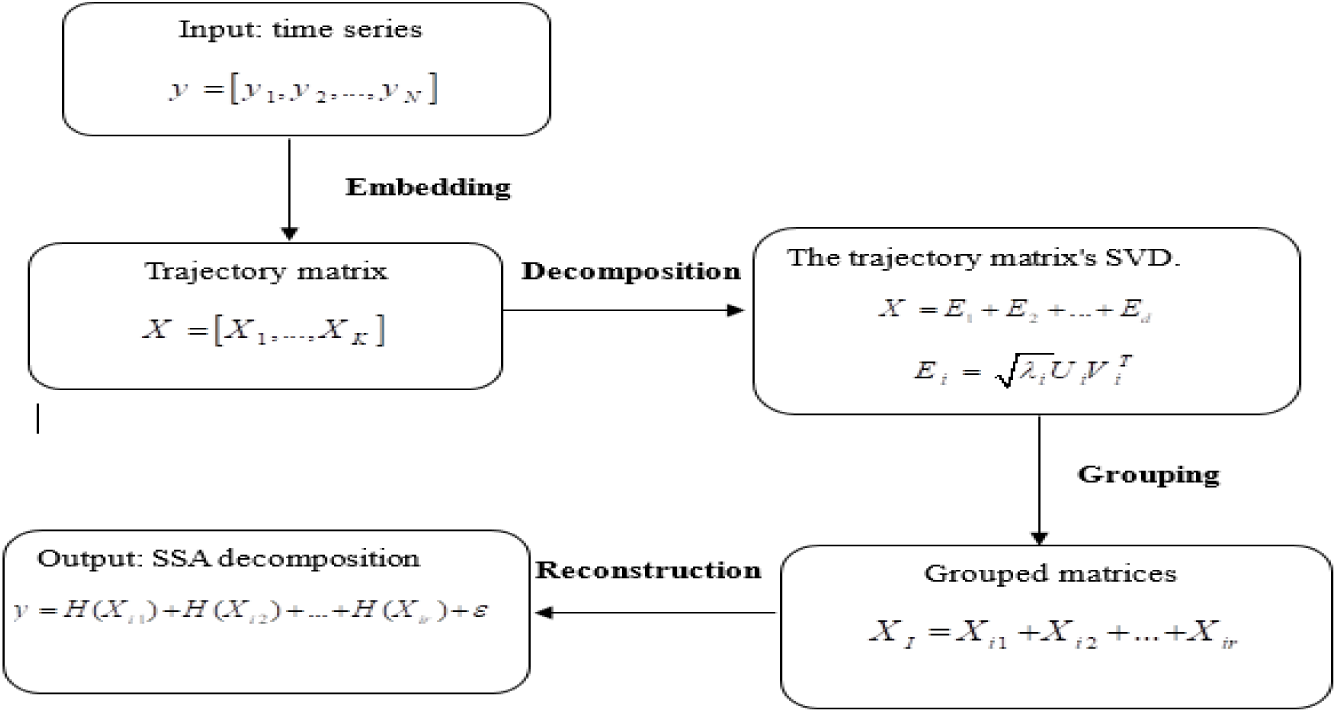
SSA family: Generic scheme.

**Figure 2.**
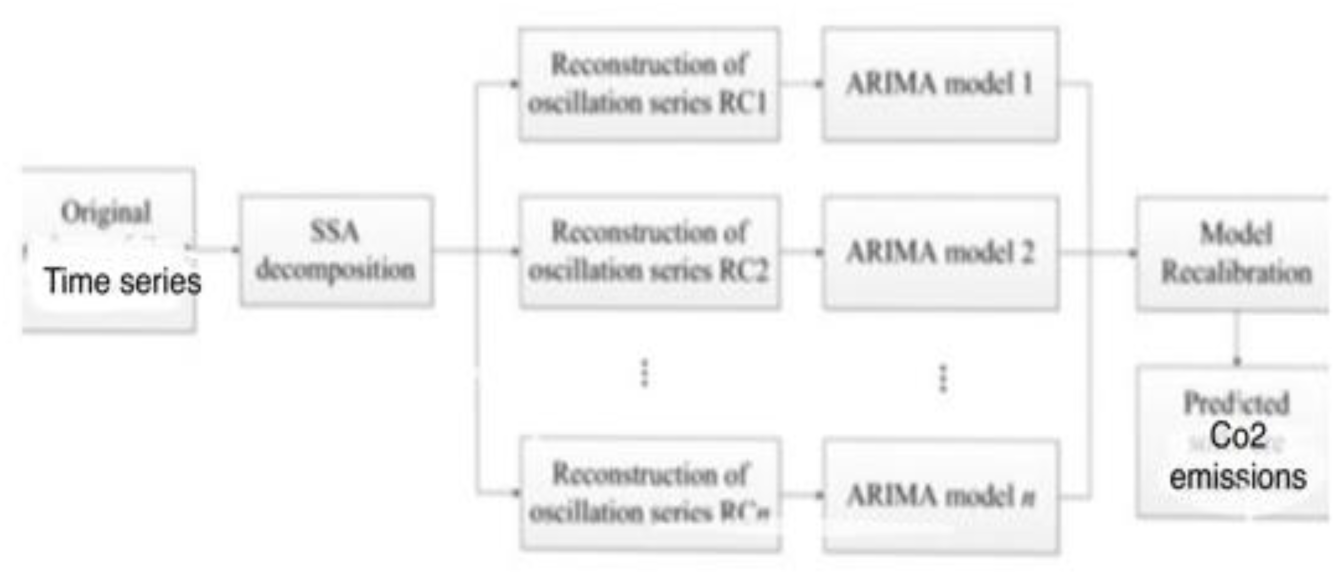
The architecture of the SSA-ARIMA model.

### 3.4 Model Performance Assessment

In this study, there were two performance evaluation criteria applied. They are calculated as shown in the next section.

#### Mean Absolute Percentage Error (MAPE)

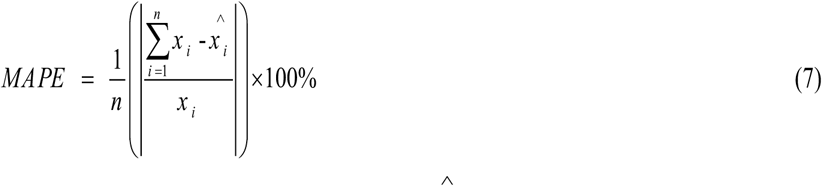

MAPE refers to mean absolute percentage error, 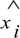 represents the predicted value, *^xi^*represents the actual value, and *n* is the total number of data observations.

#### Root Mean Square Error (RMSE)

Various statistical techniques are used to assess the estimated results in order to determine forecasting accuracy: Error Root Mean Square (RMSE).

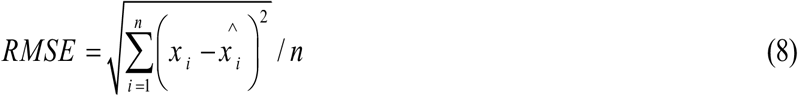

The obtained MAPE and RMSE should be as small as possible for an accurate forecast (Davalos *et al*., 2005). As indicated in Table 1, (Lewis, 1982) has provided a criterion for MAPE.

**Table.1.** Criteria of MAPE.

| Criterion (%) | Forecasting power |
| --- | --- |
| >50 | Inaccurate |
| 20-50 | Reasonable |
| 10-20 | Good |
| <10 | Highly accurate |

Source: Lewis, (1982)

## 4 Empirical Results

Three annual CO_2_ emissions (kt) in BH were used as the basis for this study’s analysis. The World Bank’s online database, which is renowned for its trustworthiness and integrity globally, provided all the data used in this analysis. Figure 3 show the graph of annual time series data for CO_2_ emissions in BH from 1990 – 2018, 2000-2018 and 2003-2018, respectively.

**Figure 3.**
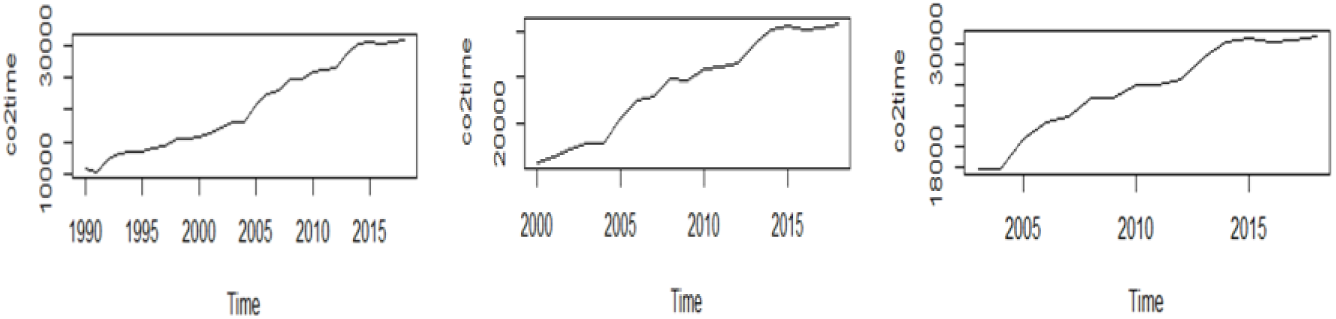
Graph of annual time series data for CO_2_ emissions in BH in three periods

### 4.1 Analysis of Singular Spectrum Analysis (SSA)

In this section, the application of the Singular Spectrum Analysis (SSA) method was applied to the time series formed by the annual time series data for CO_2_ emissions in BH to demonstrate the capability of the SSA technique to extract trend, oscillation, noise, and forecasting. This method involves two different stages such as decomposition and reconstruction. The original data series was decomposed first so that the component of the sum can be identified whether there is trend, periodic or quasi-periodic component or noise. Then, the step of reconstruction of the original time series data was done before proceeding to the forecasting of time series data.

In order to analyse the performance of the different forecasting method, the annual time series data from 1990-2018, 2000-2018 and 2003-2018 of CO_2_ emissions for BH was divided into two part which are training data and the forecasting data. There are 24, 19 and 16 of time series data used as a training data while 5 data are used for forecasting purposes for each period, respectively. The R package (Rssa) and the programme Matlab were both used to produce all of the results and graphics in the application that follows.

#### Training and Forecasting of CO_2_ emissions in BH

##### Deciding the window length

The first step to be considered is to set a window length, L with condition 1 < *L* < *N*. In this case of study, the values of L should be large as possible and smaller or equal to *N*/2.

##### Calculation of eigenvalues and eigenvectors

Let’s have a look at the time series of CO_2_ emissions in BH that was collected as set data stored in the vector X, also known as the time series’ start data. The length of the CO_2_ emissions data series in BH is N = 24, 19, 16 (training data). The time series graph shown in Figure 4.

**Figure 4.**
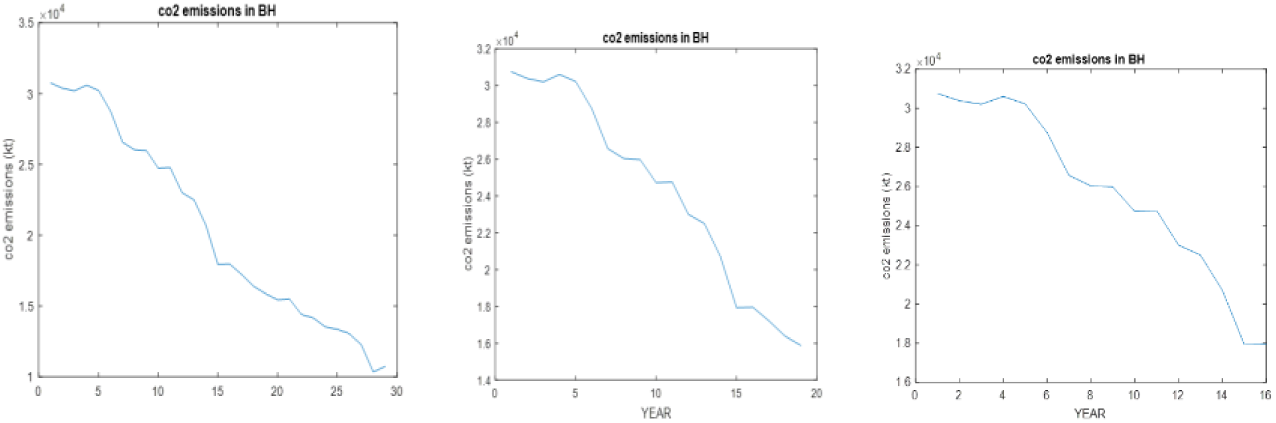
The time series of CO_2_ Emissions for BH

Since L is the window length deciding from previous step, then let K = N – L + 1. By forming K = N – L + 1 lagged vectors of size L, then mapping the initial time series data into a sequence of window length L to perform the embedding stage. The trajectory matrix S of the time series X (t) known as Hankel matrix, shows that all the elements along the diagonal are equal. Figure 5 shows trajectory matrix of the time series (*t*).

**Figure 5.**
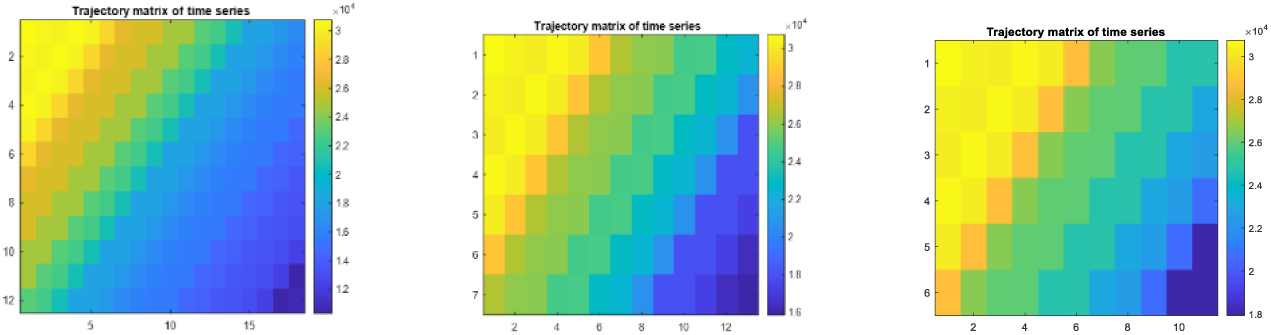
The trajectory matrix of the time series (*t*) for BH.

Following that, the trajectory matrix was subjected to singular value decomposition (SVD). This procedure is used to obtain the matrix S’s set of eigenvalues and eigenvectors. Figures 6 and 7 show the matrix S’s first and second eigenvectors and eigenvalues plotted, respectively, while Figure 8 shows a w-correlation matrix with a window length of L=12,7,6.

**Figure 6.**
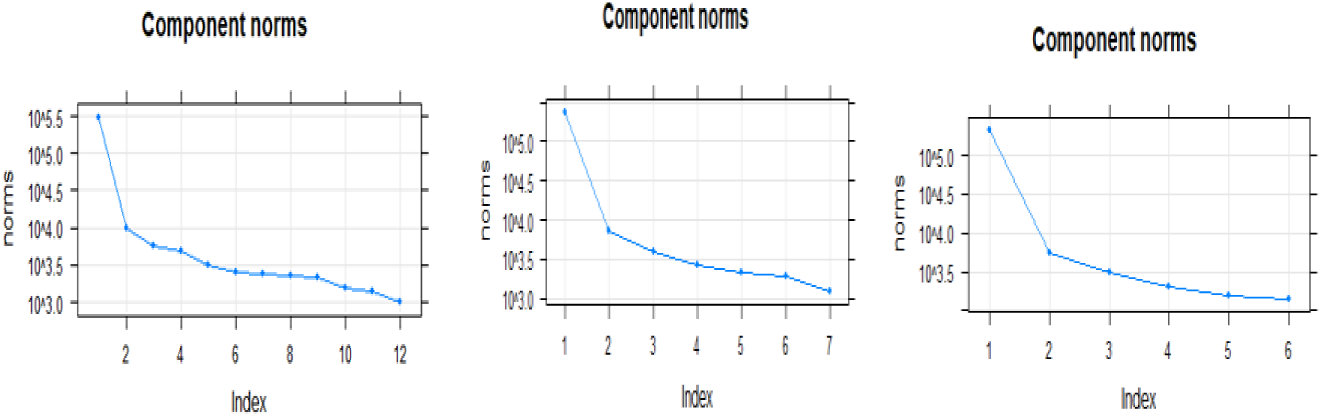
The eigenvalues of the matrix S in BH

**Figure 7.**
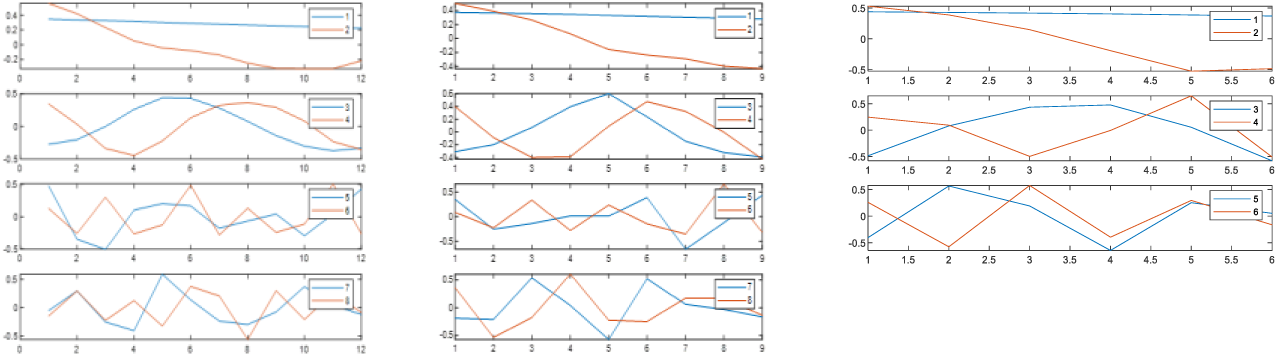
The first eigenvectors of the matrix S in BH

**Figure 8.**
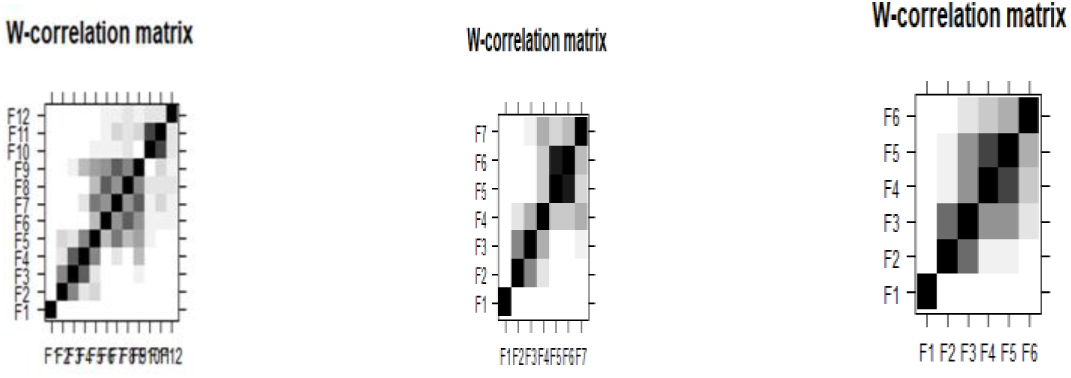
The w-correlation matrix for the 12, 7, 6 reconstructed components in BH

For appropriate identification of the sought sine waves, the graph of eigenvalues, scatterplots of eigenvectors and w-correlation matrix of the elementary components was used. The eigenvalues in Figure 8 are plotted in logarithmic scale of the 12, 7, 6 singular values for the annual time series of CO_2_ emissions in BH. Therefore, the contribution of the first few eigenvalues to the norm of X is much higher than the contribution of the rest. In fact, 99.9992% of the norm of X is contributed by the first eigenvalue, whereas the rest are much lower and vary slightly. This fact usually indicates that these last eigenvalues represent noise, or at least a nonperiodic component without any latent structure (Golyandina et al., 2001). So, several steps produced by approximately equal eigenvalues were indicated. Each step is likely to be yielded by a pair of eigenvectors which correspond to a sine wave. Since the corresponding pair of eigenvectors are similar to regular polygons. While in Figure 9 shows that the first eigenvectors is slowly-varying in BH and therefore the eigentriple (ET) should be included in the trend group.

**Figure 9.**
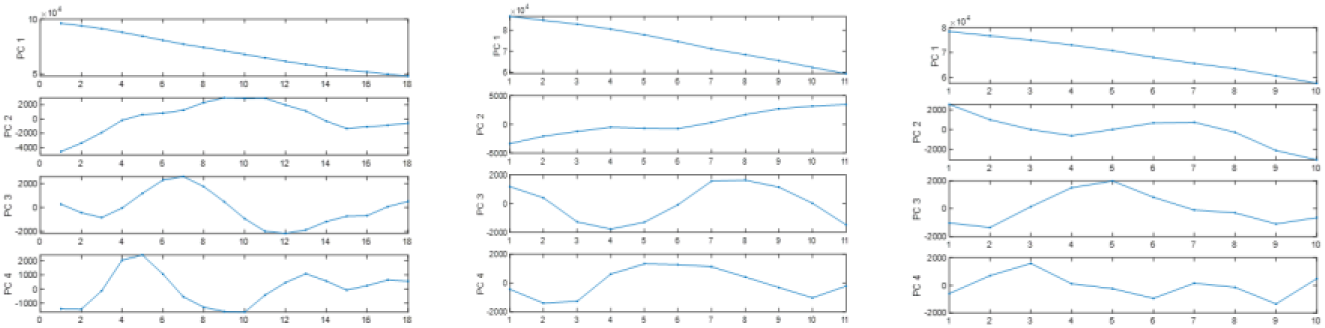
First four Principal Components of the time series.

The matrix of absolute values of w-correlations in Figure 10 is depicted in gray scale (white color corresponds to zero values, while black color corresponds to the absolute values equal to 1). This confirms that the indicated pairs are separated between themselves and also from the trend component, since the correlations between the pairs are small. While correlations between the components from one pair are very large.

**Figure 10.**
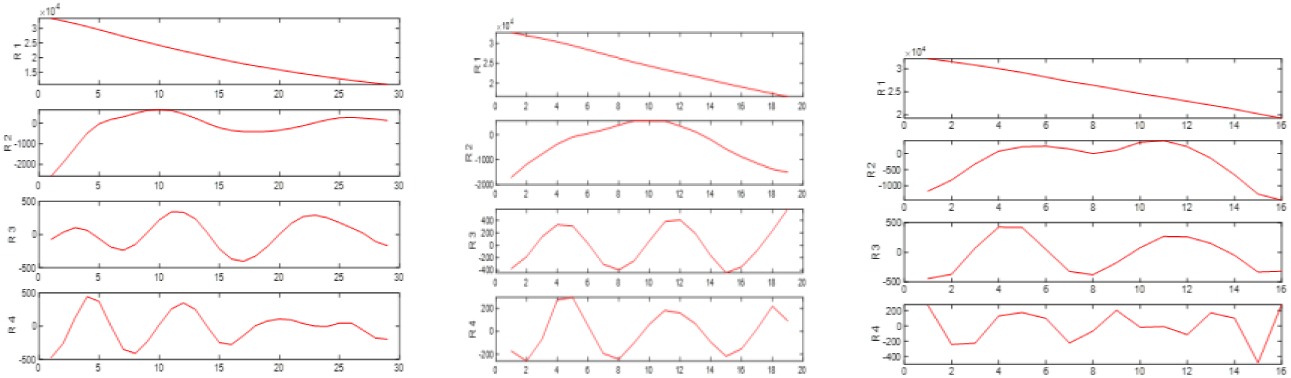
The First four reconstructed components of the time series in BH.

##### Principal components

The so-named major components of the time series X(t) are significant in SSA theory.

A pair of almost equal SSA eigenvalues and associated PCs that are roughly in phase quadrature define an oscillation. The primary components of such a pair, which are once more time series of the same length as the trajectory matrix, can indicate a nonlinear, harmonic fluctuation (Ghil *et al*., 2002).

In Matlab, the trajectory matrix X and the matrix of eigenvectors V (each column is an eigenvector) are combined linearly to calculate the principal components, resulting in a matrix of size (*N* − *L* + 1) ⨯ *L*.

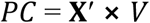

The columns of matrix *PC* are principal components of the original time series. To refer this formula we can say that “the trajectory matrix is projected onto the eigenvectors”. Figure 9 shows the first four Principal Components of the time series.

##### Reconstruction of time series

The (*N* − *L* + 1) ⨯ *L* matrix was first produced by inverting the prediction of the PC and the matrix of eigenvectors V in order to identify the rebuilt components. The reconstructed components for the initial time series are then obtained by averaging along the antidiagonals of this matrix. The reconstructed components contain a matrix size of (*N* ⨯ *L*). Figure 10 illustrates the first four reconstructed components of the time series.

According to the Figure 10 the first two reconstructed components contain practically all trends of the time series.

The comparison between the original time series and the sum of N reconstructed components is shown in Figure 11 along with the comparison between the original time series and the sum of the first two reconstructed components. It is obvious that original time series are produced by the addition of N reconstructed components.

**Figure 11.**
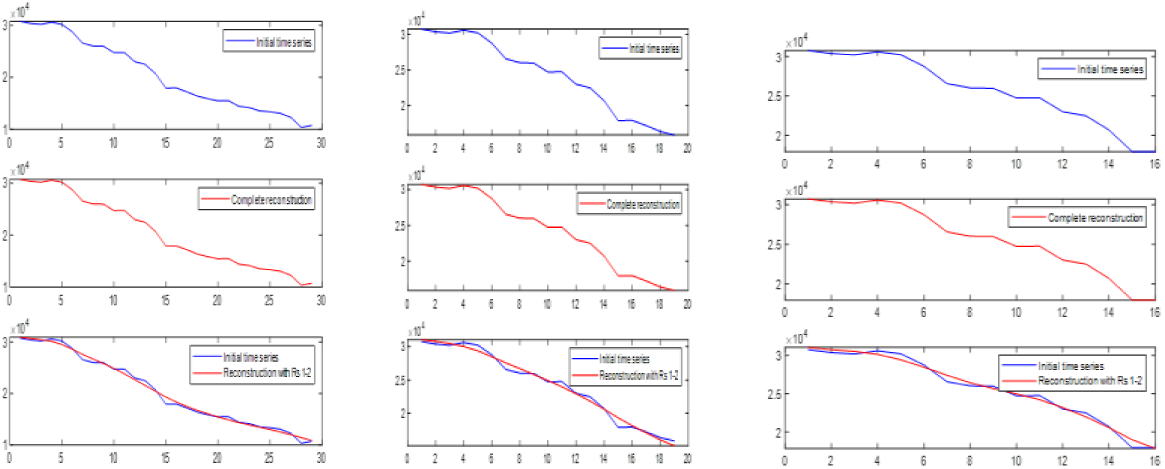
Comparison original time series and reconstructed components for BH

### 4.2 Forecasting Method

The objective of this step is to predict CO_2_ emissions for BH by applying the forecasting algorithm. Figure 12 illustrates the graph of the forecast for CO_2_ emissions of BH with window length, L=12, 7, 6.

**Figure 12.**
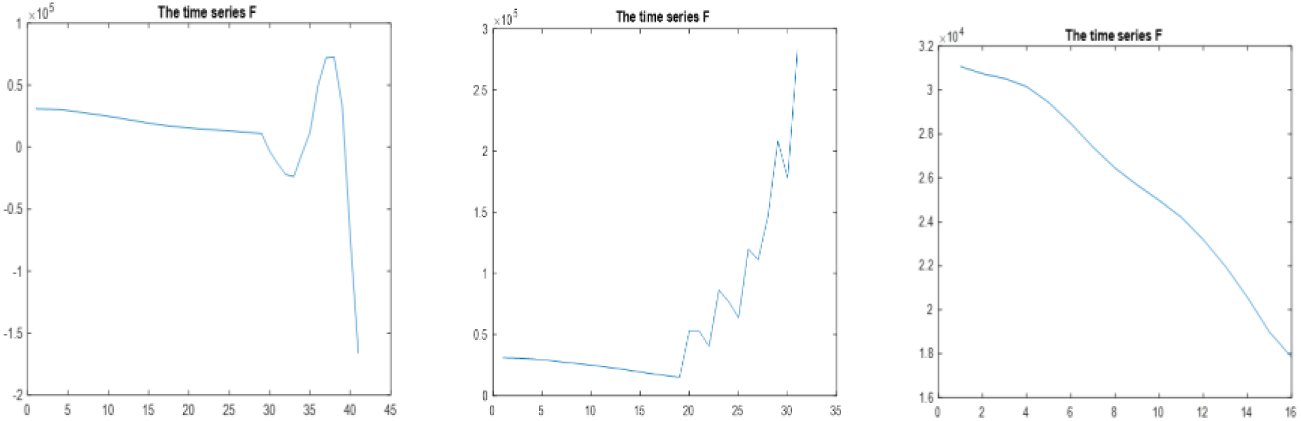
The graph of the forecast for CO_2_ emissions of BH with window length, L=12, 7, 6 for three previous periods.

It is clear from the graph that CO_2_ emission in BH is continuously increasing in the forecast values, as well as the original values (initial time series) in Figure 4. This suggests that BH will still have to deal with issues like climate change, global warming, and maintaining a clean and healthy environment.

### 4.3 ARIMA Model

To examine the stationarity of CO_2_ emissions, Augmented Dickey-Fuller1 (ADF) test (1981) was used. According to Table 2, the results of the ADF test of the time Series are not stationary in the level at which the calculated statistical significance levels are greater than the level of 0.05. The test results indicated that the time series has reached the stage of stationary after making its first difference. As indicated, the test’s statistical significance is less than the 0.05 level.

**Table 2.** Augmented Dickey-Fuller test (ADF)

| year | Critical value of ADF | The test statistic |
| --- | --- | --- |
| 1990-2018 | -4.0822 | 0.02014 |
| 2000-2018 | -3.6184 | 0.04869 |
| 2003-2018 | -7.3066 | 0.0121 |

ARIMA (0, 1, 0) for all data with lower AIC is preferable than the one with a higher AIC values (Nyoni, 2018). As a result, the ARIMA model is selected as the best as shown in Table 3.

**Table 3.** Comparison of the variants of the ARIMA models.

| Year | Box-Jenkins Model ARIMA(p,d,q) | AIC |
| --- | --- | --- |
| 1990-2018 | ARIMA(0,1,0) | 456.59 |
| 2000-2018 |  | 297.54 |
| 2003-2018 |  | 251.22 |

### 4.4 Analysis of hybrid SSA-ARIMA model

The SSA-ARIMA modelling procedure is described using the RC1 of time series for CO_2_ emissions from 1990 to 2018, 2000 to 2018, and 2003 to 2018. First, a stationary series is created using the reconstruction series RC1. The stationary behavior of the converted series’ ACF and PACF are then computed. AIC, however, establishes the appropriate order for the ARIMA model according to Table 4.

**Table 4.** ARIMA model of RC of CO_2_ emissions in BH for three periods.

| Year | RC (Sub-series) | ARIMA Model |
| --- | --- | --- |
| 1990-2018 | F1 | ARIMA 1 (2,1,1) |
|  | F2 | ARIMA 2 (2,1,3) |
|  | F3 | ARIMA 3 (0,1,0) |
|  | F (residuals) | ARIMA RES (4,1,0) |
| 2000-2018 | F1 | ARIMA 1 (1,2,0) |
|  | F2 | ARIMA 2 (2,1,1) |
|  | F3 | ARIMA 3 (2,1,1) |
|  | F (residuals) | ARIMA res (0,1,0) |
| 2003-2018 | F1 | ARIMA 1 (2,2,1) |
|  | F2 | ARIMA 2 (0,1,0) |
|  | F3 | ARIMA 3 (4,1,1) |
|  | F (residuals) | ARIMA RES (1,1,0) |

In the same way, the ARIMA model 2, ARIMA model 3, and ARIMA model L for other RC2, RC3,…RC L (L= 12, 7, 6) respectively, are determined in time series for three previous periods. The computed numbers resulting from each RC model are added up to produce the expected annual time series of CO_2_ emissions in BH. However, by selecting reasonable groups for the reconstruction series RC, we will combine the subseries 12, 7, 6 RC into a few subseries.

Eigentriple grouping and diagonal averaging are two steps in the reconstruction process. The basic matrix Y should divide into a number of groups and sum the various separated matrices in each group before anything else. Then, create a new series of length N using a number of matrices. The series that the elementary grouping reconstructs will be referred to as elementary reconstructed series. If grouping makes sense, we get a decomposition that makes sense into distinguishable series components. Typical decompositions of the results are trend, seasonality, and noise.

Checking for breaks in the eigenvalue spectra provides additional information that is used to determine the effective decomposition components. The roots of all 12, 7, 6 single values of CO_2_ emissions are shown in Fig 8 for the annual time series of CO_2_ emissions in BH of three periods.

Figure 8 demonstrates that there is a drop at component 6, which may be seen as the beginning of the noise floor that combines the feature of RCs. The leading six RCs are noteworthy, according to the results of the decomposition of CO_2_ emissions.

The characteristics of RCs are taken into account to develop suitable reconstruction components. The steps in the procedures are as follows:

1. RC1 has a distinct trend.
2. Periods of the rest RCs are computed by periodogram analysis.
3. It is important to ensure sufficient separability of the RCs when choosing window length L and recombining them. Therefore, RCs suggest that they may need to be divided into three groups in order to be reasonable, as shown in the diagram below.

The function of RSSA (in the R program) is a reconstruction that implements the reconstruction stage.

The basic signature of the function call is:
\# Reconstruction stage
\# The results are the reconstructed series r$F1, r$F2, and r$F3
recon <-reconstruct(s, groups = list(c(1,4), c(2,3), c(5,6))) res<-residuals(recon)
s: is an ssa object holding the decomposition.

Groups is a list of numerical vectors with naming capabilities that contains the indices of the fundamental reconstruction components. Based on three groups; trend, seasonal, and noise, we selected three for reconstruction and residuals (recon).

The result of the function reconstruct is simultaneously a list with the elements F1, F2, F3, and residuals (recon) that contain the reconstructed series, as well as the reconstruction object. This list can be conveniently plotted to view the reconstruction’s outcome as shown in Figure 12, and combined with an ARIMA model.

#### Model Recalibration

It also has to check to see if the superposed residual is a pure random series. Having computed the autocorrelation and the partial autocorrelation of residual error are illustrated in Figs. 13, 14, and 15, which show the annual time series of CO_2_ emissions for three different periods. The two beelines in these graphs represent at a 95% confidence level, the upper and lower bounds of the correlation coefficient and partial correlation coefficient, respectively.

**Figure 13.**
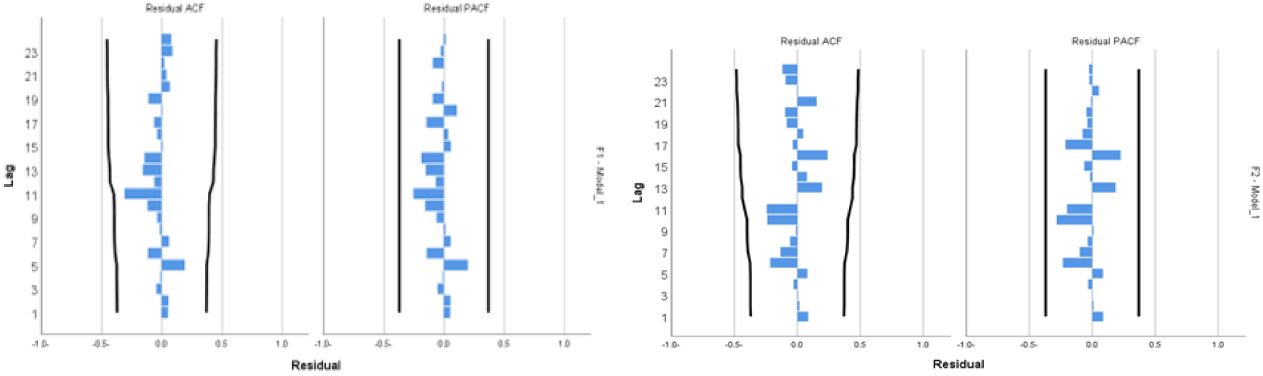
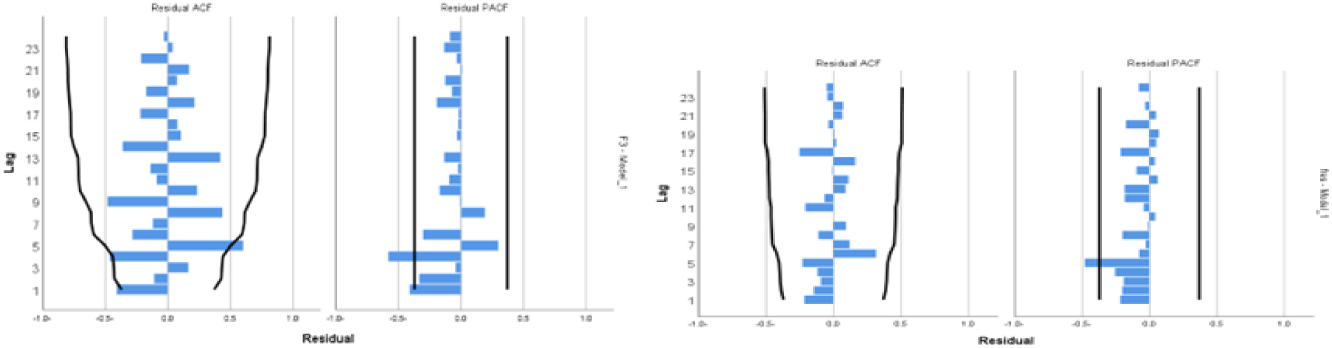
Auto correlogram and partial correlogram for CO_2_ emissions of BH in 1990-2018.

**Figure 14.**
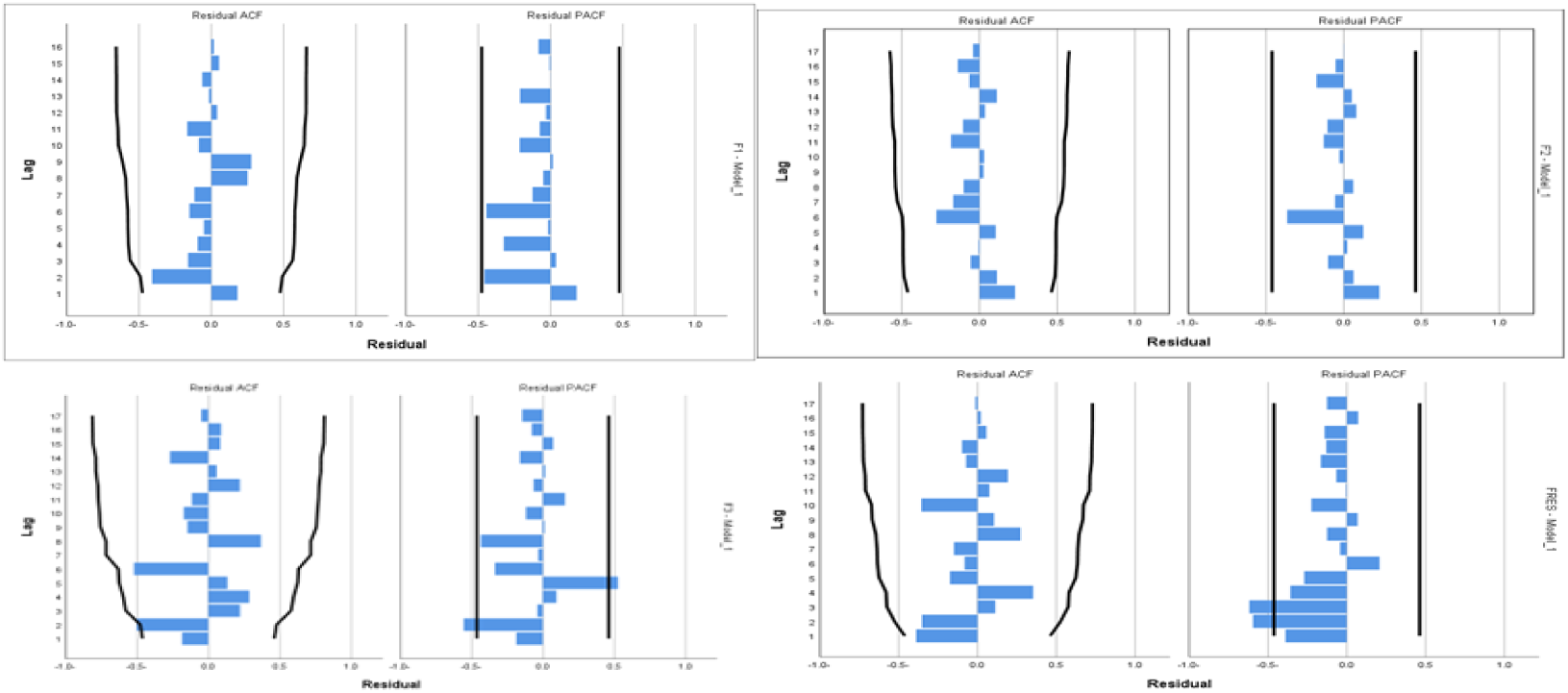
Auto correlogram and partial correlogram for CO_2_ emissions of BH in 2000-2018.

The residual error’s autocorrelation and partial autocorrelation are shown in the follow-ing figures to be negligibly small and within the confidence interval, indicating that the residual series is entirely random. As a result, the prediction result does not need to be corrected.

### 4.5 Forecasting accuracy

The forecast accuracy can be measured by the smallest Root Mean Square Error (RMSE) and Mean Absolut Percentage Error (MAPE). Table 6 below shows the comparison of MAPE and RMSE results for BH using SSA-ARIMA and ARIMA for CO_2_ emis-sions of BH.

**Table 6.** Comparison of MAPE and RMSE results for BH.

| **Table 6** Comparison of MAPE and RMSE results for BH |  |  |  |  |  |
| --- | --- | --- | --- | --- | --- |
| **TIME PERIOD** | **MODEL** | **MAPE** |  | **RMSE** |  |
|  |  | **1990-2013** | **2014-2018** | **1990-2013** | **2014-2018** |
| **1990-2018** | ARIMA(0,1,0) | 3.47% | 2.14% | 0.078039 | 0.070866 |
|  | SSA-ARIMA | 2.08% | 1.12% | 0.043096 | 0.03861 |
| **2000-2018** |  | **2000-2013** | **2014-2018** | **2000-2013** | **2014-2018** |
|  | ARIMA(0,1,0) | 3.44% | 1.69% | 0.08562 | 0.06061 |
|  | SSA-ARIMA | 1.42% | 0.91% | 0.03368 | 0.03219 |
| **2003-2018** |  | **2003-2013** | **2014-2018** | **2003-2013** | **2014-2018** |
|  | ARIMA(0,1,0) | 3.76% | 1.41% | 0.092722 | 0.055388 |
|  | SSA-ARIMA | 1.65% | 1.40% | 0.049706 | 0.0542530 |

Accordingly, Figure 16 compares CO_2_ emission predicting values in BH for all data using the hybrid SSA-ARIMA and ARIMA model. Figure 16 illustrates the significant convergence between the two models’ predicted values.

**Figure 15.**
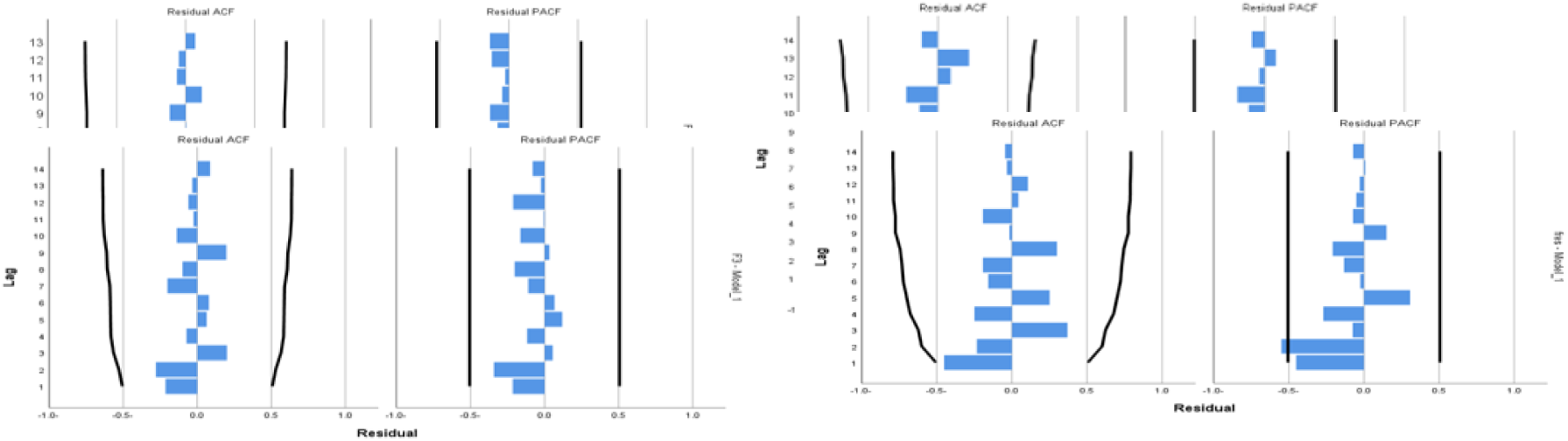
Auto correlogram and partial correlogram for CO_2_ emissions of BH in 2003-2018.

**Figure 16.**
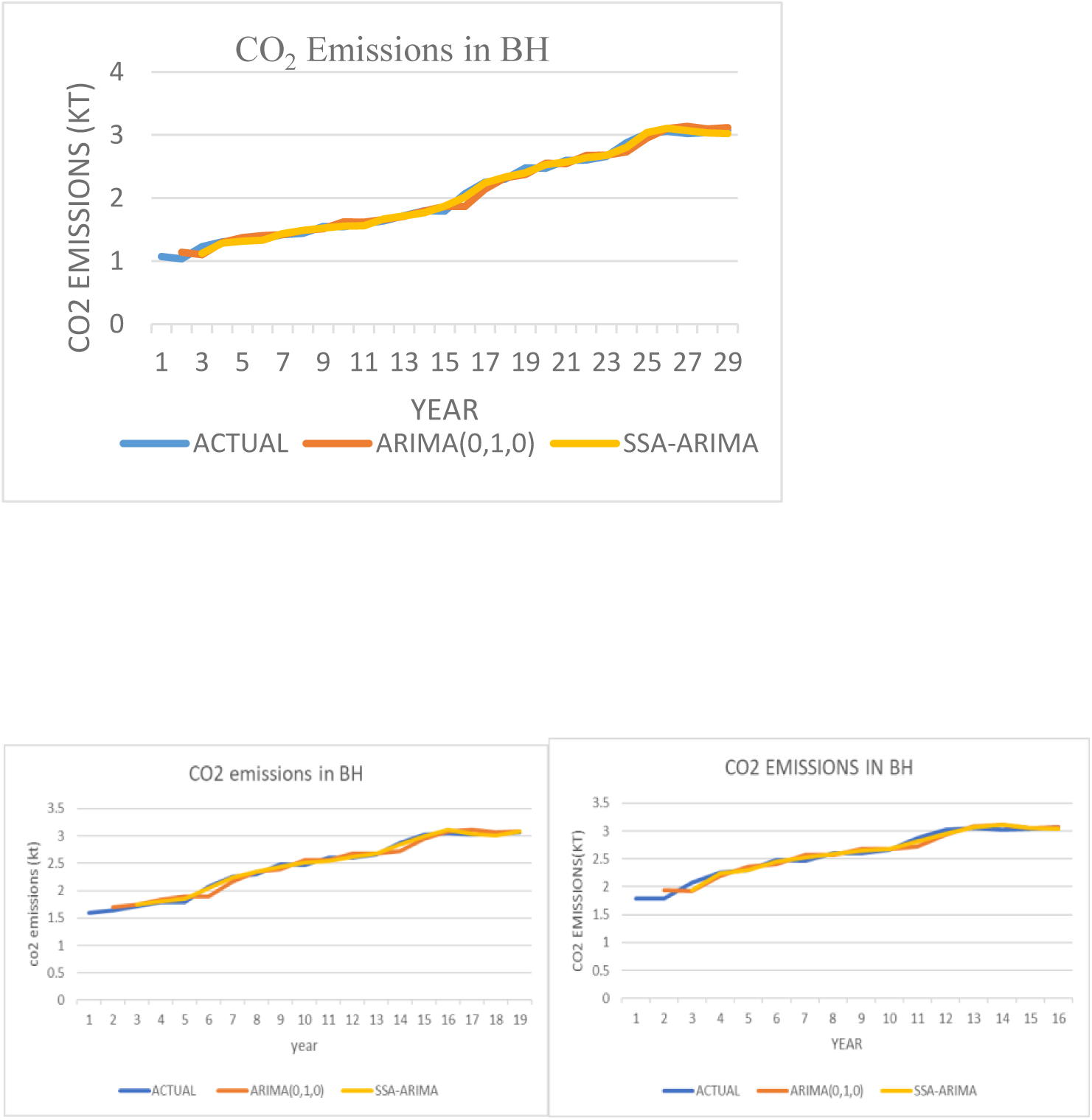
Comparison between the actual data, SSA-ARIMA and ARIMA of CO_2_ emissions in BH for all data.

In comparison to the ARIMA model, the hybrid SSA-ARIMA technique yields superior forecast accuracy. Thus, the best technique for accurately estimating CO_2_ emissions in BH is the SSA-ARIMA method. However, it is important to take into account the MAPE and RMSE values of the best model on the data from 1990 to 2018, 2000 to 2018, and 2003 to 2018.

The MAPE and RMSE values are used to assess the accuracy of forecast results for in-sample data (1990–2013), 2000–2013, and 2003–2013, as well as out-of-sample data (2014–2018). The model is more effective for forecasting if the MAPE value in the in-sample data is lower. The result of the prediction derived by the model is more accurate if the MAPE value in the out-of-sample data is lower. As a result, it can be seen that the SSA-ARIMA model performs better than the ARIMA model in both in-sample and out-of-sample MAPE and RMSE measurements.

## 5 Conclusion

This study used an empirical investigation to assess the effectiveness of the hybrid SSA-ARIMA and ARIMA model. Additionally, the ARIMA model and the SSA-ARIMA model were evaluated. To evaluate a variety of generated models, two statistical performance evaluation metrics, MAPE and RMSE, were used.

The findings of this study showed that the SSA-ARIMA model clearly affects annual time series for predicting CO_2_ emissions in Bahrain. The hybrid model’s forecasting performance is vastly superior to that of the ARIMA model. In order to show the applicability of the suggested hybrid forecasting model, future research will forecast a wider variety of time series, compare it with a wider variety of other models, and employ a wider range of metrics to assess the quality of prediction outcomes.

## 6 Acknowledgement

The authors would like to sincerely thank the University of Tabuk in Saudi Arabia for fully funding this research.

